# A Bayesian Solution to Count the Number of Molecules within a Diffraction Limited Spot

**DOI:** 10.1101/2024.04.18.590066

**Authors:** Alexander Hillsley, Johannes Stein, Paul W. Tillberg, David L. Stern, Jan Funke

## Abstract

We address the problem of inferring the number of independently blinking fluorescent light emitters, when only their combined intensity contributions can be observed at each timepoint. This problem occurs regularly in light microscopy of objects that are smaller than the diffraction limit, where one wishes to count the number of fluorescently labelled subunits. Our proposed solution directly models the photo-physics of the system, as well as the blinking kinetics of the fluorescent emitters as a fully differentiable hidden Markov model. Given a trace of intensity over time, our model jointly estimates the parameters of the intensity distribution per emitter, their blinking rates, as well as a posterior distribution of the total number of fluorescent emitters. We show that our model is consistently more accurate and increases the range of countable subunits by a factor of two compared to current state-of-the-art methods, which count based on autocorrelation and blinking frequency. Furthermore, we demonstrate that our model can be used to investigate the effect of blinking kinetics on counting ability, and therefore can inform experimental conditions that will maximize counting accuracy.

## 1 Introduction

Molecular counting aims to determine the absolute number of units in an complex, a quantity that is often essential to understanding the underlying biology of a system. The activation of T cells, for example, is sensitive to single ligands, and quantifying the molecular count of ligand-receptor interactions elucidates system sensitivity and mechanisms of signalling pathways [1]. Furthermore, molecular counting enables identification of oligomeric states, allowing differentiation of monomers, dimers, trimers, and higher order oligomers. Several biological processes depend on oligomer quantity: TGF-*β* signaling, for example, depends on the oligomeric state of Smad [2, 3]. Similarly, the oligomeric state of G-protein coupled receptors influences GPCR signaling [4, 5].

Units of interest are often separated by only a few nanometers and thus cannot be quantified through standard fluorescence microscopy techniques. While traditional fluorescence microscopy is diffraction limited to a resolution of approximately 200 nm, superresolution techniques [6, 7, 8] are able to discern individual subunits up to a resolution of 10 nm apart [9]. Below that threshold, it is no longer possible to quantify the molecular count through visual separation.

We present an alternative method of molecular counting that does not rely on super-resolution microscopy. We attempt to estimate the molecular count directly through observation of fluorescence dynamics, rather than by trying to optically separate signals. We developed a Bayesian solution to estimate the number of fluorescently labelled subunits directly from a diffraction limited spot. Our solution is based on a probabilistic model that incorporates the photo-physics of blinking fluorescent emitters and that models their dynamics over time as a Markov chain. Given a temporal record of the combined intensity of all subunits within a diffraction limited spot (Fig. 1a), this model provides the most likely number of emitters within the spot: the molecular count.

**Figure 1:**
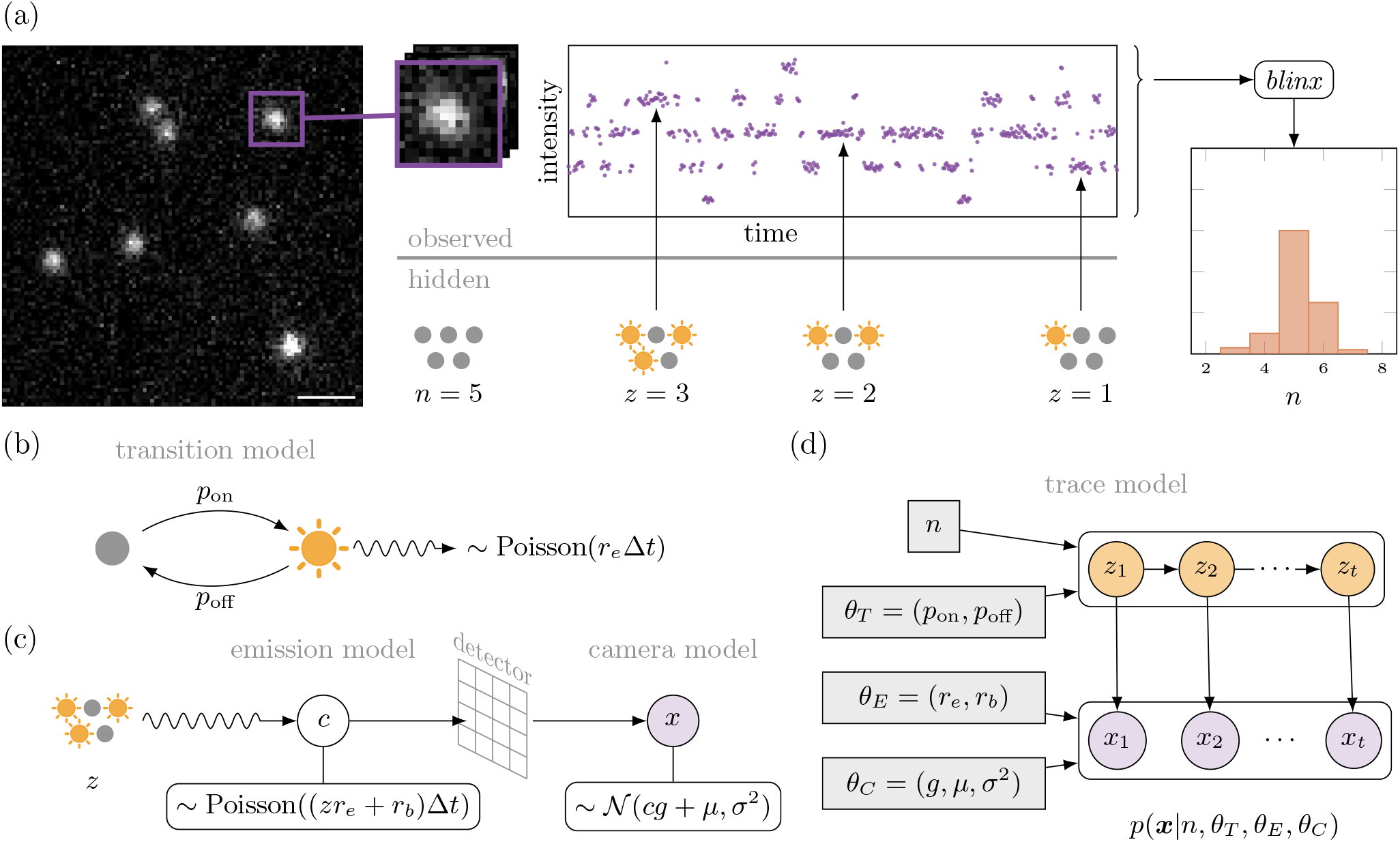
**(a)** Overview of the *blinx* method (scale bar: 1 *µ*m). *blinx* is fit on a series of observed intensity measurements, produced by a hidden state of independent stochastically blinking emitters. **(b)** *p*_on_ and *p*_off_ (*θ*_*T*_) parameterize the blinking kinetics of each individual emitter **(c)** The distribution of observed intensities is affected by both Poisson distributed shot noise (*θ*_*E*_), and Gaussian distributed readout noise (*θ*_*C*_). **(d)** *blinx* is based on a hidden Markov model conditioned on the total count *n* and seven estimated parameters.

Perhaps the simplest method of molecular counting is to correlate the combined fluorescent intensity of a spot with the number of subunits, *i*.*e*., the more subunits located within a spot, the brighter the spot is expected to be [10, 11]. This method works well for qualitative measurements, but, due to the noise in intensities measured from any single fluorophore, this approach lacks the precision required for accurate estimates of molecular counts.

Molecular counting methods that incorporate temporal fluctuations of intensity are more accurate than correlation approaches. Methods such as fluorescence correlation spectroscopy (FCS) [12, 13, 14] and balanced super-resolution optical fluctuation imaging (bSOFI) [15] fit higher order statistics to fluctuations in fluorescent intensity over time to quantify the molecular concentration. In other methods, temporal variations in intensity are induced rather than just observed; for example, counting distinct bleaching steps can provide estimates of the number of fluorescent emitters [16, 17, 18, 19].

Blinking fluorophores, such as those used in PALM [20, 21] and STORM [22], can be used to facilitate solving the counting problem [23, 24]. The blinking behavior allows for modeling at the single-molecule level, compared to the bulk measurements of FCS, while the repeated transitions in intensity provide more information than the irreversible switches that occur in bleaching-based counting. A major limitation of these methods is the complex photophysical properties of these blinking fluorophores, making it difficult to differentiate between possible dark states and photo-bleaching.

In contrast, DNA-PAINT [25] achieves blinking through transient DNA-binding, effectively decoupling blinking from photophysics. This has the additional advantage of producing blinking fluorescence that is functionally immune to photo-bleaching, due to the continuous replenishment of fluorophores from solution [26].

Quantitative DNA-PAINT (qPAINT) [27] estimates the molecular count based on the frequency of blinking events. For example, if a blinking rate of one event per second is calibrated to one molecule, an observation of ten events per second corresponds to a count of ten molecules. qPAINT is a relative counting method, and as a result is entirely dependant on the quality of the calibration, which can be a non-trivial endeavor. Localization based FCS (lbFCS+) [28, 29] combines the structured blinking of DNA-PAINT with the principles of FCS, fitting the autocorrelation function of intensity over time to produce a count. lbFCS+ can accurately count up to six molecules in a diffraction limited spot, without calibration. However, both of these methods are limited because they fit summary statistics, rather than the data directly. In contrast, we fit the model to every measurement, fully utilizing both the time and intensity information.

Furthermore, all prior methods provide a single estimate of molecular count, even though a probabilistic count may be preferable for downstream analysis. Bayesian approaches can estimate a likelihood for each possible condition and have been used previously to infer the number of FRET conformational states [30, 31], the number of photo-bleaching steps [19], the assignment of blinking events to specific fluorophores [32, 33], and the molecular count from observed intensity [24].

We propose *blinx*, a Bayesian model to estimate the molecular count directly from a trace of fluctuating diffraction limited fluorescent intensity measurements. Based on a fully differentiable hidden Markov model, *blinx* fits seven parameters that describe the photo-physics and kinetics of the system, to the sequence of measurements and estimates a likelihood for each possible count. These likelihoods can be compared directly, producing a posterior distribution of counts given the observed sequence. We first run *blinx* as a forward model, to simulate different experimental conditions and to determine the effect of each condition on counting ability. We find that *blinx* provides a substantial improvement in calibration-free (compared to lbFCS+) and calibrated (compared to qPAINT), counting accuracy, and can count across different kinetic regimes. Finally, we prove the counting ability of *blinx* experimentally by validating the estimated count with ground truth measured by super-resolution DNA-PAINT imaging.

## 2 Method

Starting from the photo-physics of the system, we derive a probabilistic model that accounts for both the observed intensity and the temporal fluctuation kinetics. Given an intensity trace, the output of our method is the posterior distribution over molecular counts, providing not only the most likely count, but also a comparison to all other possible counts.

### 2.1 Model

We consider the case of multiple fluorescent emitters within a single diffraction limited spot. As such, we can measure only their combined intensity contributions *x*_*t*_ at time *t*. We assume that each emitter blinks stochastically and independently of the others, yielding a fluctuating trace ***x*** = (*x*_1_, …, *x*_*T*_) of total intensity measurements over *T* frames (Fig. 1a). Our goal is to infer the number of emitters *n*, given ***x***.

Formally, we are modelling a posterior distribution of the number of emitters *n* given the intensity trace ***x***, *i*.*e*.:

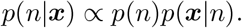

Assuming a uniform prior on the number of emitters, *p*(*n*) can be ignored and it remains to model the likelihood *p*(***x***|*n*). We model this likelihood as a hidden Markov model where the hidden state, denoted by *z*_*t*_, represents the number of emitters that are active and emitting photons at time *t* (see Fig. 1d). The hidden Markov model is conditioned on a set of parameters *θ*, including *θ*_*C*_ camera parameters, *θ*_*E*_ emission parameters, and *θ*_*T*_ transition parameters,

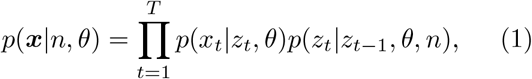

where we define *p*(*z*_1_|*z*_0_, *θ, n*) = *p*(*z*|*θ, n*), *i*.*e*., the initial distribution of the hidden state *z* is the stationary distribution.

Following this framework, the likelihood is the product of two probability distributions: an intensity model *p*(*x*_*t*_|*z*_*t*_, *θ*) and a transition model *p*(*z*_*t*_|*z*_*t*−1_, *θ, n*).

#### 2.1.1 Intensity Model

The distribution of observed intensity *x*_*t*_ given the number of currently active emitters *z*_*t*_ can be directly derived from the photo-physics of our microscope. In the following section we omit the subscript *t* for clarity.

Working backwards through the light path, the measured intensity value *x* is a function of the number of photons *c* detected (Fig. 1c). The detector itself contributes noise to the system, known as the readout noise, which we model as *p*(*x* |*c, θ*_*C*_). The number of photons hitting the detector, in turn, depends on the number of active emitters *z* and is distributed following a shot noise model *p*(*c*|*z, θ*_*E*_).

The probability of observing an intensity *x* given *z* can then be obtained by marginalizing over *c, i*.*e*.:

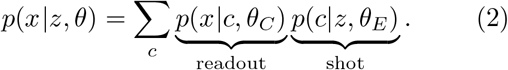

##### Readout Noise

The readout noise from the detector is often assumed to be negligible (*i*.*e*. when using an EMCCD camera). However, when using an sCMOS camera, its contribution is significant [34]. To accommodate both systems, here we include the readout noise in our model. However, the readout noise contribution can be negated by setting *µ* = 0, and *σ*^2^ << 1. Given *c* photons, the readout intensity of the camera is normally distributed [34]

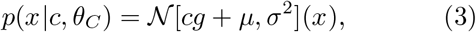

where *θ*_*C*_ = (*g, µ, σ*^2^) are the camera’s gain, offset, and variance, respectively.

In the following, it will be beneficial to express the readout noise as a zero mean normal distribution that is independent of the number of photons detected. To that end, we transform our measurement *x* into photon space by accounting for the camera gain and offset:

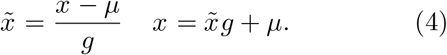

We can now express our readout noise distribution in terms of the transformed measurement 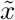, which we further shift by −*c* to obtain a zero-mean distribution:

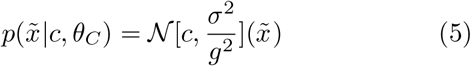

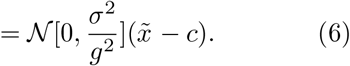

##### Shot Noise

Due to shot noise, the number of photons detected over a time interval Δ*t* is Poisson distributed [35] (Fig. 1c). Our model accounts for two sources of emitted photons: first, the *z* active emitters produce photons at a rate *zr*_*e*_, and second, out of plane emitters produce a relatively constant number of background photons *r*_*b*_. Combined, these sources produce the expected number of photons *λ* = (*zr*_*e*_ + *r*_*b*_)Δ*t* per frame.

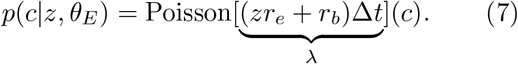

For large values of *λ*, the Poisson distribution approaches a normal distribution with both mean and variance of *λ* and therefore we approximate our shot noise distribution as:

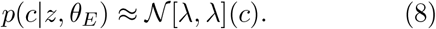

##### Combined Intensity Model

Taking together the readout noise transform (Eq. 6) and shot noise approximation (Eq. 8), the intensity distribution (Eq. 2) can now be rewritten as

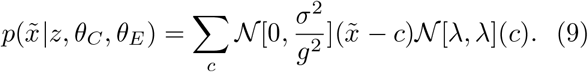

Similar to [34], we note that this expression resembles a convolution, *i*.*e*., the sum of two independent random variables (the photon count *c* and a zero-mean, photon-independent camera readout noise, (Eq. 6)). Since the two distributions are normal, we can rewrite the above as a single normal distribution with summed means and variances:

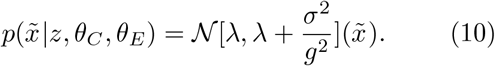

#### 2.1.2 Transition Model

To model the step-like temporal fluctuations in intensity observed in the trace ***x***, a distribution is needed that describes the change in the number of active emitters *z* over time. To accomplish this, we assume that the process is Markovian and the number of active emitters *z*_*t*_ at time *t* depends only on the number of active emitters at the previous time point *z*_*t*−1_. On the individual emitter level, we define *p*_on_ as the probability of an inactive emitter at time *t* − 1 to become active at time *t*. Conversely, we define *p*_off_ as the probability of an active emitter at *t*− 1 to become inactive at time *t* (Fig. 1b).

For the case of a single emitter (*i*.*e*., *n* = 1), the probability of observing this emitter as active (*i*.*e*., *z*_*t*_ = 1) if the emitter was inactive in the previous frame, is *p*_on_, or (1 − *p*_off_) if it was previously active (and similarly for the probability of observing the emitter as inactive). For multiple emitters, however, a transition from *z*_*t*−1_ to *z*_*t*_ can be caused by multiple events. Consider, for example the case of two emitters (*n* = 2). A transition from *z*_*t*−1_ = 1 to *z*_*t*_ = 1 can be caused either by no change occurring or by one emitter becoming active while the other emitter deactivates.

In the general case (*n* ≥ 1), we consider all possible events that can lead to an observed change in *z*. To that end, we marginalize over the number *a* of active emitters that deactivated from *t* − 1 to *t*:

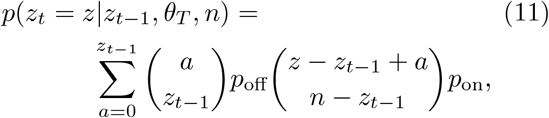

We assume that all emitters share the same *p*_on_ and *p*_off_, and that both probabilities remain constant over time.

#### 2.1.3 Inference

The likelihood *p*(***x***|*θ, n*) of observing ***x***, see (Eq. 1), depends on a total of seven parameters *θ* and the total number of emitters *n*. However, we are interested in the posterior distribution *p*(***x***|*n*). Accounting for the priors on *θ* in addition to the likelihood, this is known as the model evidence and is defined as:

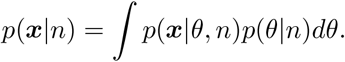

Although no closed form solution exists for this integral, multiple approximations exist including variational methods [31] and the Laplace approximation [36]. We use the latter, which is the maximum likelihood solution corrected by the Occam factor:

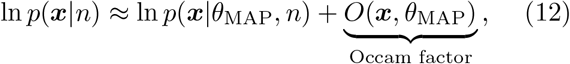

where the Occam factor *O*(***x***, *θ*_MAP_) is defined as

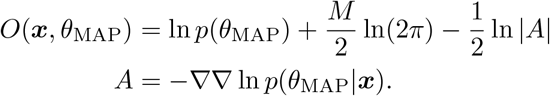

Here, *M* is the dimensionality of *θ*, which is constant. Because we are interested only in comparing likelihoods, this term can safely be ignored, resulting in:

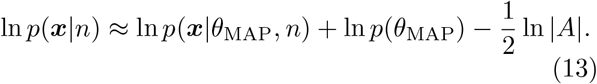

Therefore, to estimate the posterior distribution over *n*, we first need to find the maximum likelihood parameters *θ*_MAP_ for each value of *n*.

This model is fully differentiable, allowing us to estimate the maximum likelihood parameters through gradient ascent:

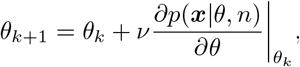

where *ν* is the step size, dynamically calculated by the Adam optimizer, and the gradients of *θ* are taken with respect to the likelihood: *p*(***x***|*n, θ*). The maximum likelihood parameters *θ*_MAP_ are then obtained,

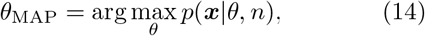

thus allowing us to estimate the posterior distribution *p*(***x***|*n*). Additionally, we can obtain an estimate of the maximum likelihood molecular count:

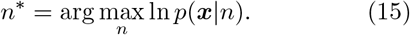

## 3 Results

### 3.1 Simulated Experiments

Running *blinx* as a forward model to simulate experiments, we generated traces for counts *n* = 1 to 30 (Fig. 2a,b). We determined experimentally relevant camera parameters from a calibration of the sCMOS camera (*θ*_*C*_: *g* = 2.17, *µ* = 4791, *σ*^2^ = 774). We determined kinetic (*θ*_*T*_ : *p*_on_ = 0.036, *p*_off_ = 0.028), and emission parameters (*θ*_*E*_: *r*_*e*_ = 2.79, *r*_*b*_ = 6.77) from an initial experiment, and these values closely matched those reported in the literature [29]. We simulated traces for 4000 frames, corresponding to an imaging time of 14 minutes.

**Figure 2:**
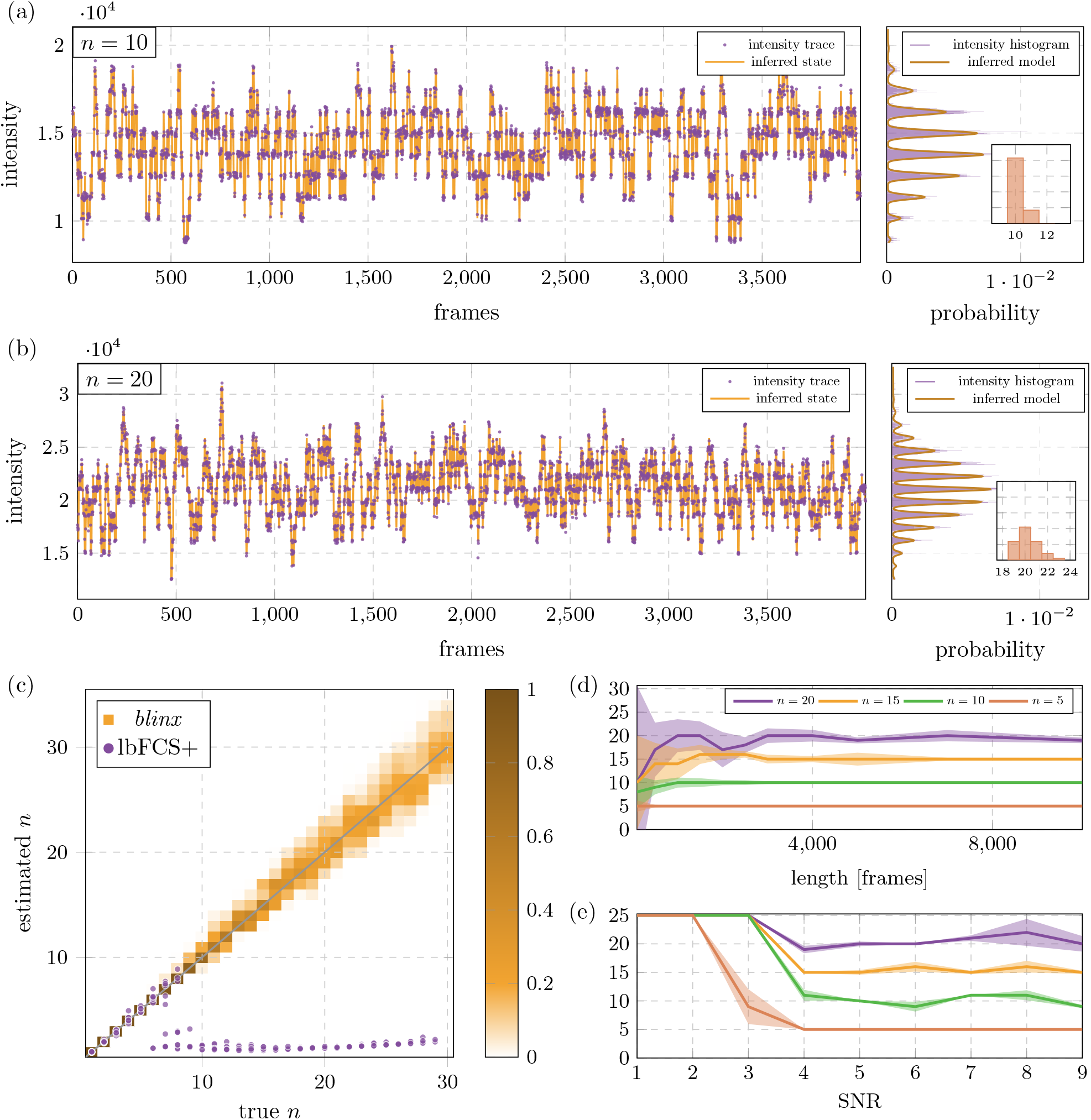
**(a, b)** Traces simulated from *n* = 10 (a) or 20 (b) emitters, and a density plot of measured intensities. The inferred state and inferred model demonstrate accurate fitting of both the intensity and transition distributions. Inset: posterior distribution estimated by *blinx*. **(c)** Estimated versus true *n*, shown as posteriors for *blinx* (orange heatmap) and point estimates for lbFCS+ (purple dots). **(d)** As trace length increases, the variance of the *blinx* posterior decreases, with limited improvement above 4000 frames. **(e)** At low signal-to-noise, *blinx* incorrectly estimates the maximum count tested as the most likely.

For inference, we placed tight, empirically determined, Gaussian priors on the camera parameters.

We also placed flat uniform priors on the kinetic parameters, allowing the model to fit blinking rates without external bias, and loose priors on the emission parameters (*r*_*b*_, *r*_*e*_) to prevent over-estimation of count (see Section 4). We experimentally determined the prior on *r*_*b*_ by averaging the intensity of the pixels immediately outside the region of interest. We estimated the prior on *r*_*e*_ by first fitting a count of *n* = 1 to the trace. The distribution of all observed *z*-states is unimodal. Therefore, whichever *z*-state is the most common, the second most common state will be *z* ± 1. We observed that fitting *n* = 1 captures this specific transition, and provides a good estimate of photon emission rate of a single emitter *r*_*e*_.

*blinx* accurately fit simulated time series with counts up to *n* = 30 (Fig. 2c). For *n* = 10 and below, *blinx* estimated the true count with a substantially higher likelihood than all other possibilities. Above *n* = 10 the posterior broadened as other possibilities became more likely. The most likely estimate was the true count in 177/300 (59%) of traces and was within 1 of the true count in 267/300 (89%) of all traces. Importantly, in many cases *blinx* is able to fit the true count despite the fact that not every state was occupied; *i*.*e*. the model is not simply counting the number of peaks in the intensity histogram (Fig. 2a,b). This is a tripling in performance compared to lbFCS+, which counted accurately up to *n* = 6 and was within 1 of the true count for 64/300 (21%) of traces, but failed to estimate higher counts. Upon further analysis, the main limitation of lbFCS+ is the estimation of the intensity of a single emitter, corresponding to *r*_*e*_ in *blinx*. lbFCS+ relies on a histogram based method to determine this value, while *blinx* is able to jointly optimize this parameter with other parameters. Providing this value to lbFCS+ led to a substantial recovery of performance. With *r*_*e*_ provided, lbFCS+ identified 200/300 (67%) of traces within 1 of the true count.

Both *blinx* and lbFCS+ count without prior knowledge of the kinetic parameters. In contrast, qPAINT is heavily dependant on a kinetic calibration and therefore is not comparable in this context. A direct comparison to qPAINT is shown in Section 3.3.

Longer time series provide more information to the model than shorter series and increase counting accuracy. We simulated and fit time series of lengths ranging from 100 to 10,000 frames, with the same parameters as the previous section (Fig. 2d). As expected, accuracy increased and variance of the posterior distribution decreased, as length increased. Interestingly for short time series, *blinx* consistently underestimated the true count. This is likely because only a subset of possible states were observed in this short time.

We determined the robustness of our model and the minimum signal-to-noise ratio (SNR) needed to achieve an accurate count. We artificially increased the variance of the readout noise *σ*^2^, to simulate traces with SNRs ranging from 9 to 1 (where 9 corresponds to the parameters previously used). Because noise in our model is a function of intensity, quantifying the SNR of a trace is not trivial. For simplicity, here we define SNR as the difference in intensity between the first two states (*z*_0_ and *z*_1_) divided by the standard deviation of the difference. In effect this is the higher bound of SNR for a given trace. *Blinx* shows accurate counting for SNRs ≥ 4 (Fig. 2e). Interestingly, for SNR ≤ 4, *blinx* estimates the count as 25, the highest *n* tested, no matter the true count.

This is due to the model adding states to compensate for the wide distribution of intensities (see Section 4).

### 3.2 Effect of kinetic parameters

The performance of *blinx* depends on the true kinetic parameters of the system (*p*_on_ and *p*_off_). To find a regime that maximizes the accuracy of *blinx*, we simulated and fit traces from *n* = 1 to 20, with a range of kinetic parameters, while holding all other parameters constant. The parameter values included the range of experimentally determined kinetics: qPAINT: (*p*_on_, *p*_off_) = (0.006, 0.2) [27] and lbFCS+ (0.02, 0.02) [29]. We summarized the resulting posteriors by calculating the expected squared error over all true counts: Σ*w*(*n*^*∗*^−*n*)^2^, where *w* is the probability of the estimated count *n* (Fig. 3a).

**Figure 3:**
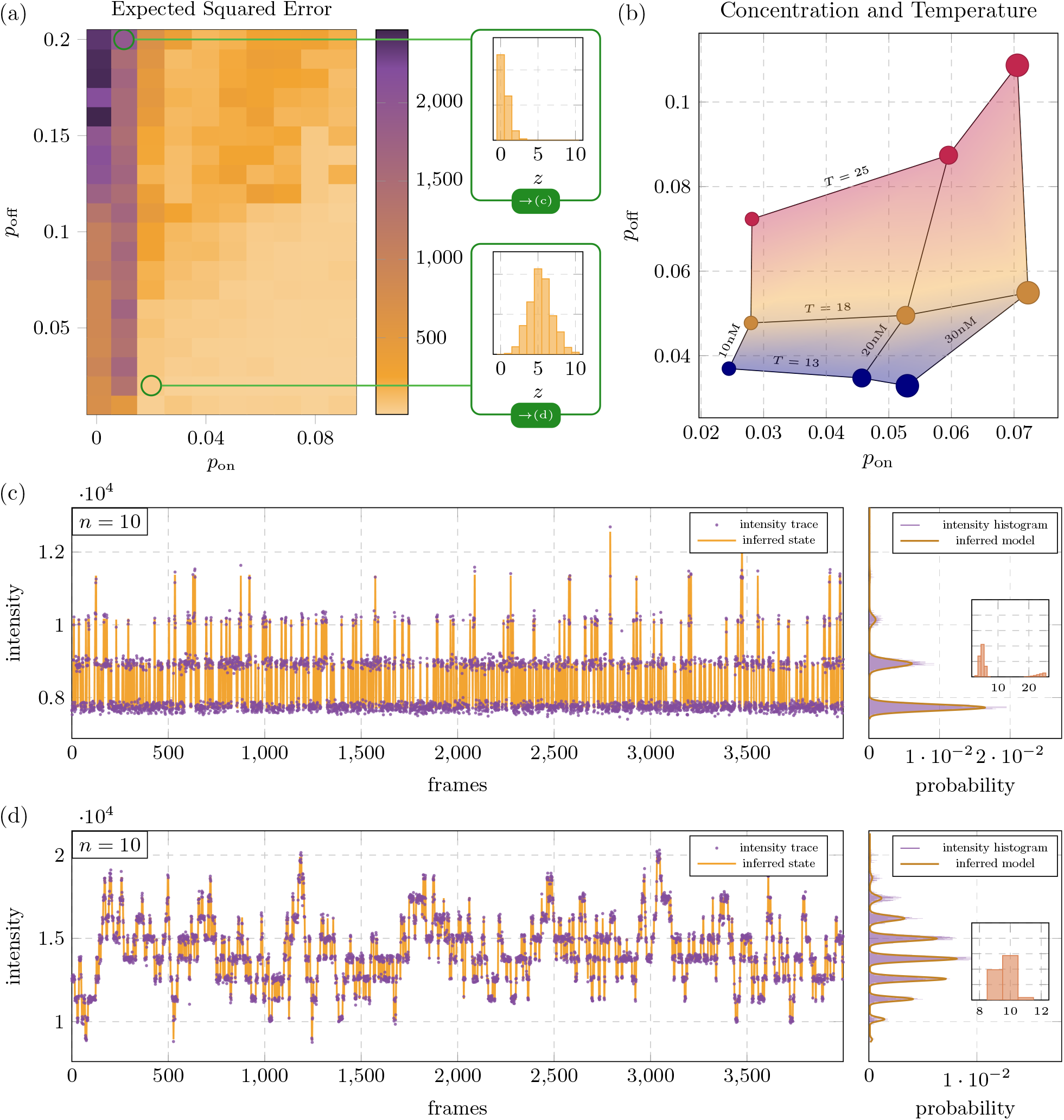
**(a)** The counting ability of *blinx* is dependant on the blinking kinetics, showing increased model accuracy when *p*_on_ > *p*_off_. Top inset: when *p*_on_ < *p*_off_, the distribution of observed *z* states is shifted towards 0. Bottom inset: for *p*_on_ = *p*_off_, the distribution of *z* states is centered at 1*/*2 *n*. **(b)** Average experimental blinking kinetics as a function of imager concentration (marker sizes) and temperature (color). **(c)** Trace and *blinx* fit, *p*_on_ =0.01, *p*_off_ =0.2, and true count of *n*=10, representative of (a) top inset. **(d)** Trace and *blinx* fit, *p*_on_ =0.02, *p*_off_ =0.02, and true count of *n*=10, representative of (a) bottom inset.

The results can be divided into two regimes: *p*_on_ < *p*_off_ and *p*_on_ ≥ *p*_off_. In the first regime, where *p*_on_ < *p*_off_, we see significant limitations to our models counting ability. Furthermore, when *p*_on_ is substantially lower than *p*_off_, our model loses almost all ability to count and estimates *n* = 1 or 2, regardless of the true *n*^*∗*^. In contrast, in the second regime, where *p*_on_ ≥ *p*_off_ we observe accurate counting up to *n* = 20. A clear difference between these two regimes can be seen in the distribution of their hidden states *z* (Fig. 3a, insets). When *p*_on_ < *p*_off_, the distribution is shifted towards 0, and a majority of the frames occupy the lowest two *z*-states (Fig. 3c). In this regime, the true count becomes indistinguishable for all methods, without prior information. When *p*_on_ ≥ *p*_off_, this distribution is centered, or even shifted towards *n* and a larger fraction of the states are visited (Fig. 3d). This provides ample information for the model to infer the correct *n*.

In DNA-PAINT, the blinking rate is determined by the kinetics of single-stranded DNA binding, which in turn depends on experimental conditions such as temperature and concentration, and the specific DNA sequence [37]. As a result, the kinetic parameters *p*_on_ and *p*_off_ of our model can be tuned by adjusting the temperature and imager concentration. As seen in (Fig. 3b), *p*_on_ increases with imager concentration and *p*_off_ increases with temperature. We imaged DNA origami with a known count of *n* = 1 at 25°C and an imager concentration of 10 nM and measured *p*_on_ = 0.028 and *p*_off_ = 0.072 (Fig. 3b), conditions within the poor counting accuracy regime of *p*_on_ < *p*_off_ (Fig. 3a). To examine performance in the predicted more favorable kinetic regime, we increased imager concentration and decreased temperature. Increasing imager concentration to 30 nM raised *p*_on_ to 0.071 and decreasing temperature to 13°C decreased *p*_off_ to 0.037 (Fig. 3b). As expected, the effects of temperature and concentration were largely independent of one another [37]. We also observed a substantial decrease in SNR with decreasing temperature and increasing concentration. This was an expected side effect of increasing concentration, because increasing imager concentration increases *r*_*b*_. But the effect of temperature on SNR was surprising. We hypothesize that this is due to the stabilization of partial binding between imager and docker strands at low temperatures. Balancing the increase in counting accuracy and the decrease in SNR, we identified imaging conditions of 20 nM and 13°C as optimal on our microscope.

### 3.3 qPAINT Kinetic Regime

qPAINT relies on the accurate measurement of the average dark time between blinking events [27] and is therefore optimized for a kinetic regime where blinking events are short and infrequent (Fig. 4a,b). This regime presents a challenge to *blinx* (Fig. 3a top inset). If the only states ever observed are *z* = 0 or 1, there is insufficient information to estimate count without prior knowledge. qPAINT faces the same limitation and relies on a calibration of the blinking kinetics of a single binding site. Because *blinx* is a Bayesian model, it can incorporate a calibration as priors of the kinetic parameters. With this addition, and the tightening of the priors on *r*_*e*_ and *r*_*b*_, the counting accuracy of *blinx* is restored (Fig. 4c). *blinx* estimated the true count as the most likely in 133/300 (44%) of traces and estimated within 1 of the true count in 269/300 (90%) of all traces, comparable to the favorable kinetic regime (Fig. 2c). However, obtaining these priors requires additional experimental steps, and is often not trivial. Thus, a tradeoff exists. Without any calibration, *blinx* is able to accurately count in a subset of kinetic conditions where *p*_on_ > *p*_off_. But with experimental calibration, the counting ability of *blinx* is expanded to a wider range of kinetic conditions. Further, counting in this kinetic regime expands the use of *blinx* beyond diffraction limited spots, and enables counting in super-resolved SMLM reconstructed images.

**Figure 4:**
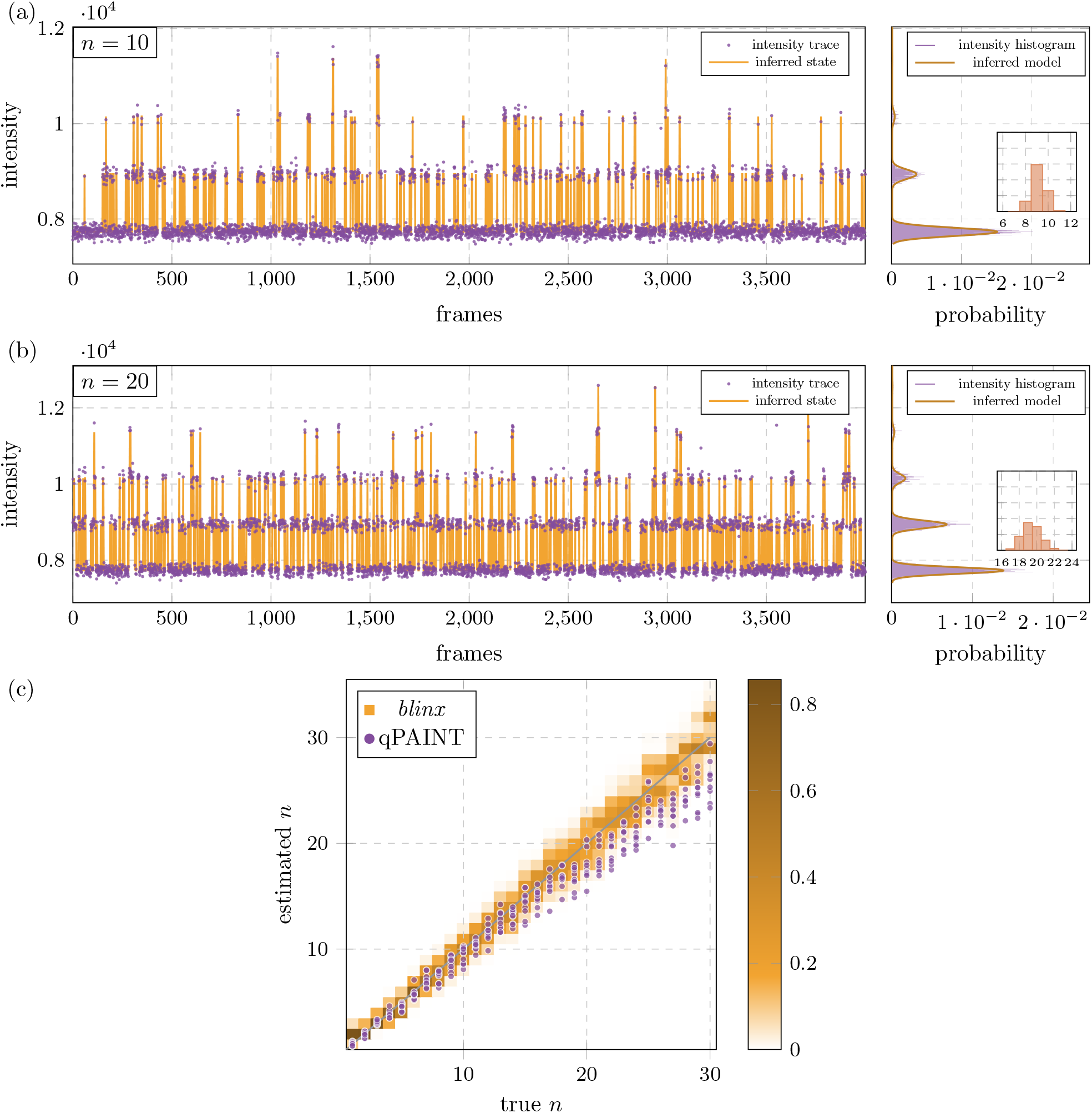
**(a, b)** Simulated traces and *blinx* fits, *p*_on_ =0.006, *p*_off_ =0.2, and *n* = 10/20 emitters. *blinx* was fit with strong priors on all parameters. **(c)** Above *n* > 15 qPAINT underestimates the count, while the posterior of *blinx* remains accurate on average.

Due to the stochastic nature of blinking, multiple emitters can be active at any given time, which becomes increasingly likely at higher counts. This is not compatible with the qPAINT requirement of well separated, single emitter blinking events. As a result, at higher *n* the measured average dark time is longer than the true average dark time and results in qPAINT underestimating molecular count at higher *n* (especially noticeable above *n* = 20, see Fig. 4c). In *blinx*, the simultaneous activity of multiple emitters is modeled explicitly in both the intensity and transition models ((Eq. 7) and (Eq. 11)). As a result, *blinx* avoids underestimation and can provide an accurate molecular count up to *n* = 30.

### 3.4 Experimental Counting

To experimentally validate the performance of *blinx*, we used DNA-Origami, which provides nano-scale control over the number and location of emitters [38]. We designed DNA-Origamis containing 1 and 4 DNA-PAINT docker strands, spaced in a grid 20 nm apart. This distance was chosen so that the true number of docker strands could be visually confirmed through super-resolution post-processing. Incorporation efficiency is roughly 80 percent for each docking site [39], so only a fraction of the origamis were expected to contain all 4 dockers. Origamis were first imaged at 13°C with low laser power (1.5%) and 20 nM imager concentration to collect traces (4,000 frames at 200 ms exposure time) for counting with *blinx* (Fig. 5a,b). The system was then brought to 25°C, buffer exchange was performed, and new imager was added at 10 nM. The origamis were imaged again at high laser power (40%), and post-processed with Picasso [25], to obtain super-resolution ground truth (Fig. 5c). Only origamis that had a visual count matching the designed count (1 or 4) were selected for analysis with *blinx*. Of the 131 traces with a known count of 1, *blinx* correctly counted 112 (85%) and the average of the posterior distributions is shown in Fig. 5d. Possible counts up to *n* = 8 were modelled, but the likelihoods for counts above *n* = 3 were negligible for all traces.

**Figure 5:**
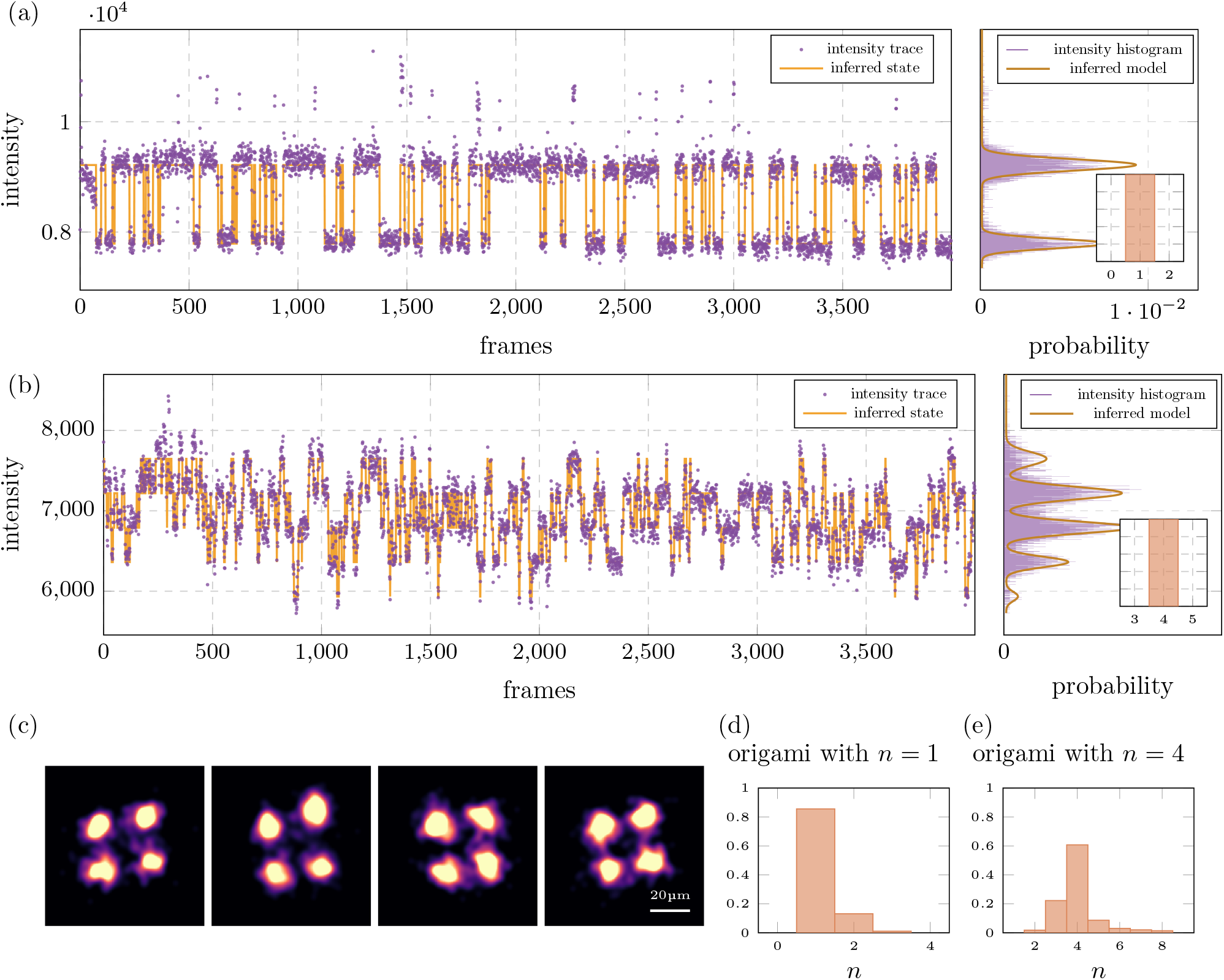
**(a, b)** Experimental traces with super-resolution confirmed counts of one (a) and four (b), and *blinx* fits **(c)** Superresolution visualization of origamis made with four binding sites. **(d, e)** Average *blinx* posterior distributions of 131 traces with a known *n* of one (d) and 110 traces with a known *n* of four (e)

For the traces with a known count of 4, 71/110 (65%) were correctly identified as 4, and 103/110 (94%) were identified as between 3 and 5. The average posterior distribution is shown in Fig. 5e. Importantly, traces were not preprocessed or filtered before undergoing *blinx* counting analysis.

## 4 Discussion

Our analysis of different kinetic regimes highlights a trade-off between resolution and counting. Relatively fast kinetics are required to generate temporal separation required for single molecule localization microscopy. In contrast, a slower kinetic regime, where multiple emitters are active in a single frame, is optimal for counting. *blinx* can accurately estimate molecular count in both regimes, either calibration-free in the slow regime or with a calibration in the fast regime.

While *blinx* is robust to a range of experimental conditions, we identified several limitations. First, *blinx* tends to overestimate the molecular count in time series with low SNR. The model expects intensities to be observed in evenly spaced, normally distributed peaks (see histograms in Fig. 5a,b). With low SNR, noise can generate intensity between peaks, and adding additional *z*-states can sometimes yield higher likelihoods than correctly absorbing this noise into the fitted distribution. In the limit, an infinite number of states could perfectly explain any intensity time series. This can be avoided by specifying a prior on *r*_*e*_, the photon emission rate of a single fluorophore, which effectively increases the cost of adding additional states, and therefore decreases the likelihood of these models.

A second limitation is that the intensity model does not capture all of the noise in the system. As seen in Fig. 5a, there are occasional frames where the measured intensity is greater than expected and fits the model distribution poorly. To compensate for these outliers, we incorporated a baseline outlier probability into the intensity model, *i*.*e*., a fixed uniform likelihood for observing any intensity within the recorded range of values. To further reduce the influence of these observed outliers on counting accuracy, we excluded the highest 0.5% of intensities values when calculating the likelihood (the specific value is adjustable as a hyperparameter).

In this work, we used DNA-PAINT to test the performance of *blinx* due to its resistance to photobleaching and consistent, predictable blinking behavior. However, *blinx* is not specific to DNA-PAINT and could be applied to any experimental system that meets the following three assumptions.

First, all sub-units are assumed to behave independently. In the intensity model, this means that observed intensity will scale linearly with the number of active emitters *z*. In the transition model, this means that the blinking kinetics of one spot have no effect on the blinking kinetics of any other. In practice, subunits in close proximity might interact and violate this assumption [40]. Other methods, like qPAINT, also make this assumption and have been shown to work well in experimental systems [41, 42].

Second, all sub-units within a spot are assumed to have identical properties (*p*_on_, *p*_off_, *r*_*e*_). Rather than model each sub-unit individually, we model the summed behavior of all sub-units combined. In other words, we assume that each sub-unit is interchangeable with any other within the same spot. In some applications, where non-uniform behavior among subunits may occur, *e*.*g*., if one unit is less accessible to diffusion of imager than the others [43]. In these cases, we expect a decrease in model performance. Finally, all properties are assumed to remain constant over time. Experimentally, drift in emission properties (*r*_*e*_, *r*_*b*_) can result from an unstable focus and should be corrected before any processing with *blinx*. Changes in kinetic properties over time are also sometimes observed and could be caused by photo-bleaching, damage to emitters or temperature changes. Unlike the first two assumptions, this assumption is not fundamental to the structure of *blinx* and slight extensions of our model could account for these time dependant parameters. In fact, we anticipate that intentional temporal changes in parameters could be used to gain more information from the system and further increase model performance.

In principle, these assumptions could be relaxed within the Bayesian framework of *blinx*, providing opportunities to apply *blinx* to new systems and questions. For example, the simple modification of setting *p*_on_ = 0 could support the counting of photobleaching events. Additionally, by incorporating dynamic priors, *blinx* could also capture changing conditions over time, potentially increasing counting ability beyond what we demonstrate here. Further, with minor modifications to the transition distribution to account for photobleaching, this method could be extended to other stochastically blinking emitters, such as those used in PALM or STORM, opening the door for molecular counting in living samples.

## 5 Code Availability

Code for the model *blinx* is available at https://github.com/funkelab/blinx.

Code and data for the experiments described can be found at https://github.com/funkelab/blinx_experiments.

## 6 Experimental Methods

### DNA Origami

DNA origami with 20 nm spaced docking strands were designed with the Picasso design module [25] and assembled following the procedure in [25], with single-strand DNA oligos purchased from IDT. A repetitive 11 nt docker sequence (CTCCTCCTCCT) was added to the 3’ end of select staples as the docker sequence. Such repetitive sequences have been shown to increase *p*_on_ [43]. DNA origami samples were prepared for imaging in Ibidi *µ*-Slide 8-well glass bottom slides, again following the procedure described in [25].

### DNA-PAINT

DNA-PAINT imaging was performed following the standard procedure detailed in [25]. Imaging solution contained buffer B (5 mM Tris-HCl, 10 mM MgCl_2_ 1 mM EDTA, pH 8) and an oxygen scavenging system consisting of PCA, PCD, and Trolox. Super-resolution images were acquired with a 7 nt imager (GAGGAGG-Cy3B, Biosyn) at a concentration of 10 nM, while counting analysis images were acquired with a 8nt imager (GAGGAGGACy3B, Biosyn) at a concentration of 20 nM. All images were post-processed in the Picasso-localize module to detect spots and correct drift over the time course.

### Microscopy

TIRF imaging was performed on a Zeiss Elyra 7 microscope, equipped with a pco.edge 4.2 sCMOS camera (pixel size 6.5 *µ*m), and an *α* Plan-Apochromat 63x/1.46 TIRF oil immersion objective. An Okolab H101-Cryo-BL system provided temperature control. It was found that this temperature control system was not compatible with super-resolution experiments, due to vibrations caused by the pump that were transferred to the objective through the cooling jacket. Therefore for super-resolution experiments, the temperature control system was turned off, and the sample temperature allowed to increase to room temp (25°C) before imaging.

## 7 Acknowledgments

We thank Brian English, Chris Obara, Lorena Benedetti, and Jennifer Lippencott-Schwartz for their expertise in super-resolution microscopy and for sharing use of their microscopes. We thank James Liu for his advice on DNA-PAINT and on troubleshooting imaging systems. We thank Diane Adjavon, Caroline Malin-Mayor, Manan Lalit, Magdalena Schneider, and Cedric Allier for their help in model development and python implementation of the model. We thank Shalin Mehta for insight into modeling and calibrating the photo-physics of these systems. We thank Mark Aronson for his help reviewing and editing this manuscript. We further thank Ralf Jungmann for helpful comments on our approach. J.S. acknowledges support from the European Molecular Biology Organization (ALTF 816-2021). We thank Petra Schwille and Florian Stehr for sharing previously published raw data. We thank George M. Church and Chao-ting Wu for hosting A.H. for an initial training visit. This work was supported by the Howard Hughes Medical Institute and the Janelia Visiting Scientist Program.

## 8 Author Contributions

A.H. and J.F. derived the model and wrote all code. A.H. performed all simulations and microscopy experiments. J.S. provided training on DNA-PAINT and DNA-Origami experiments and analysis. P.T. and D.S. provided valuable insights and guidance for both method development and experiments. A.H. and J.F. wrote the manuscript. A.H., J.F., P.T. and D.S., conceived the project. All authors revised the manuscript and have given approval to the final version.

## Notes

### Competing Interest Statement

The authors have declared no competing interest.

### Summary of Updates

Fixed minor typos in the text. No changes to any data or figures

